# Gene drive-based population suppression in the malaria vector *Anopheles stephensi*

**DOI:** 10.1101/2024.05.24.595689

**Authors:** Xuejiao Xu, Jingheng Chen, You Wang, Yiran Liu, Yongjie Zhang, Jie Yang, Xiaozhen Yang, Zhengbo He, Jackson Champer

## Abstract

Gene drives are alleles that can bias the inheritance of specific traits in target populations for the purpose of modification or suppression. Here, we constructed a homing suppression drive in the major urban malaria vector *Anopheles stephensi* targeting the female-specific exon of *doublesex*, incorporating two gRNAs and a *nanos-*Cas9 promoter to reduce functional resistance and improve female heterozygote fitness. Our result showed that the drive was recessive sterile in both females and males, with various intersex phenotypes in drive homozygotes. Both male and female drive heterozygotes showed only moderate drive conversion, indicating that the *nos* promoter has lower activity in *A. stephensi* than in *Anopheles gambiae*. By amplicon sequencing, we detected a very low level of resistance allele formation. Combination of the homing suppression and a *vasa*-Cas9 line demonstrated a boost in the drive conversion rate of the homing drive to 100%, suggesting the use of similar systems for population suppression in a continuous release strategy with a lower release rate than SIT or fRIDL techniques. This study contributes valuable insights to the development of more efficient and environmentally friendly pest control tools aimed at disrupting disease transmission.

## Introduction

Vector-bone diseases, including malaria, Dengue fever, and West Nile Virus, continue to pose a global health threat, causing numerous annual fatalities worldwide. Disease vector management is crucial for halting transmission. However, overuse of chemical-based treatments has accelerated the emergence of pesticide resistance^1^. In the ongoing effort to combat these diseases, various genetic engineering tools, such as irradiation/chemosterilant-induced or transgene-based sterile insect technique (SIT)^2–4^, release of insects carrying a dominant lethal (RIDL)^5–7^, and *Wolbachia*-mediated incompatible insect technique (IIT)^8–10^, have been developed. Notably, gene drive technology stands out as a highly efficient population control tool with the potential to affect a whole population through a minimal release.

The concept of homing gene drive, utilizing selfish genetic elements, was proposed two decades ago^11^, but its development was hindered by lack of ability to target specific sequences. The advent of CRISPR genome editing tools marked a significant breakthrough, enabling more sophisticated and practical gene drive strategies. A homing gene drive consists of a Cas9 endonuclease and gRNA cassette, capable of cleaving the target site, generating double-strand break in wild-type allele, and utilizing homology-direct repair to copy the drive allele into the wild-type allele. This converts a heterozygote into homozygote in the germline. This process enables the drive to bias its inheritance in the offspring, so called Super-Mendelian inheritance. However, if end-joining repair takes place instead of homology-direct repair, the target site can be mutated. This is called a resistance allele because it can no longer be cleaved by Cas9/gRNA, hindering gene drive transmission.

While gene drive development has been explored not only in model species such as fly^12–14^, mice^15,16^ and *Arabidopsis*^17,18^, as well as non-model organisms like yeast^19^, herpesviruses^20^, and agriculture pests^21^, the efficiency in insect species beyond a few studies in *Drosophila*^12,13,22^, *Aedes*^23^ and especially *Anopheles* mosquitoes^24–26^ has been relatively lower due to varies factors such as low conversion rate, high fitness cost, parental effects and resistant allele formation. Several potential solutions have been proposed to address these issues. For instance, drive conversion and fitness of drive carriers can be promoted by improved regulatory elements or coding sequences^27,24,28,22^, while carefully selected targets^24,27^, “toxin-antidote” (or “Cleave and Rescue”) systems^29,12,13^ and gRNA multiplexing^12,13,30^ can help in removing resistant alleles.

Gene drive strategies primarily aim at population modification or suppression. Population suppression gene drives have garnered attention for their potential to directly remove pests by disrupting reproductive capabilities or biasing the sex ratio of a population. Common targets include essential female fertility genes. In proof-of-principle studies in *Drosophila melanogaster* targeting *yellow-G*, only moderate drive equilibrium frequency was achieved in cage populations^30^, limiting suppressive power. This was caused by lack of high drive conversion rates, moderate fitness costs, and moderate rates of resistance allele formation from early embryo cleavage by maternally deposited Cas9 and gRNA. Drives utilizing improved germline promoters^28^ or decoupling of drive and fertility gene cleavage^14^ could mitigate these issues. Another promising target for population suppression is the sex determination pathway, with the female-specific exon of *doublesex* (*dsx*) emerging as a key target. Successful suppression in laboratory cage trials was achieved in *Anopheles gambiae*. However, functional resistance and fitness costs remain as potential problems in larger populations^24,31^. A different interpretation of previous experimental data^32^ indicated that the *nanos* promoter could be have equal or better performance for drive conversion rate and especially fitness costs, which inspired our design in this study.

Although studies on modification drives have been published in *A. stephensi*, the major urban malaria vector in Asia that is becoming more invasive in East Africa, these lack efficiency and/or suitable cargo genes to be release candidates. Also, no suppression gene drive study has been reported for this species. Here, we constructed a suppression drive (named HSDdsx) as well as a *vasa*-Cas9 line. The suppression drive exhibited intermediate drive inheritance rate and minimal resistance, indicating success of the 2-gRNA design but failure of the *nanos-Cas9* allele to achieve high cut rates. Crosses of both lines resulted in significantly improved inheritance rate (to 100%) in HSDdsx, suggesting their use in a continuous-release deployment for substantially more efficient population suppression compared to SIT and fRIDL. Our study lays the groundwork for the further construction of highly efficient gene drives for population management of *A. stephensi* and provides valuable insights for other non-model organisms.

## Methods

### Plasmid design and construction

Donor and helper plasmids were generated employing the Gibson assembly method. The donor plasmid designed for homing-based suppression, denoted as HSDdsx, contained a Cas9 coding sequence controlled by the *nanos* promoter, a 3xP3-EGFP-SV40 fluorescence marker, two distinct gRNA cassettes under the control of the U6a or 7SK promoters, and flanking homology arms facilitating homology-directed repair-mediated integration. The U6a-gRNA target (gRNA1: 5’-ttcaactacaggtcaagcgg-3’) was strategically positioned at the highly conserved intron 4-exon 5 boundary, consistent with a prior *Anopheles gambiae* study^24^. The target of 7SK-gRNA (gRNA2: 5’-cgcaataccacccgtcagag-3’) was situated within exon 5, 56 bp downstream of gRNA1.

An independent Cas9 helper plasmid was also constructed, featuring a Cas9 coding sequence driven by the *vasa* promoter, to facilitate knock-in for both the HSDdsx and *vasa*-Cas9 constructs. Additionally, two gRNA-expressing helper plasmids were developed exclusively for the transformation of *vasa*-Cas9, which contained a recoded *hairy*, a Cas9 coding sequence driven by *vasa* promoter, a 3xP3-EGFP-SV40 fluorescence marker and a U6-gRNA cassette. The target sites of these two gRNA plasmids (KI-gRNA1: 5’-cacacatccaaaatggtgac-3’; KI-gRNA2: 5’-ggccaccagccagataccgc-3’) were located around the translation start site of *hairy*, enabling successful integration of the *vasa*-Cas9 construct and subsequent translation of the recoded coding sequence for gene function rescue.

All the regulatory elements and target gene sequences were identified by reciprocal BLAST analysis of their homologs in *D. melanogaster* against the genome of *A. stephensi* through NCBI database (https://blast.ncbi.nlm.nih.gov/Blast.cgi). gRNA target sites, their activity, and their potential off-target sites were analyzed by using the online tool CHOPCHOP (https://chopchop.cbu.uib.no/).

All plasmids were constructed using Hifi DNA Assembly Cloning Kit (NEB, USA) and then midipreped with ZymoPure Midiprep Kit (Zymo Research, USA). Plasmid sequences were confirmed with Sanger sequencing by BGI. The final plasmid sequences are available at Github (https://github.com/jchamper/ChamperLab/tree/main/Anopheles-stephensi-dsx-Drive).

### Mosquito rearing

All the mosquitoes were maintained at 27±1°C, 75% humidity, and a 12h light/dark cycle. Larvae and adults were provided with fish food (Hikari, Japan) and a 10% sucrose solution, respectively. Adult mosquitoes were housed in 30 cm x 30 cm cages for mating, and females were blood fed using the Hemotek blood-feeding system (Hemotek, UK) with defibrinated cow blood.

### Embryonic microinjection and germline transformation

Three to four days post blood meal, plastic cups covered with wet filter paper were put into the cage for a 30-minute interval to collect eggs, which were subsequently lined up for microinjection. The injection mix for generating the HSDdsx line comprised 152 ng/μL of the donor plasmid, 300 ng/μL of the *vasa*-Cas9 helper plasmid, and 300 ng/μL of Cas9 protein. For the *vasa*-Cas9 line, the injection mix included 613 ng/μl of the donor plasmid, approximately 100 ng/μL of each gRNA plasmid, and 170 ng/μL of the *vasa*-Cas9 helper plasmid. Surviving G0 mosquitoes were subsequently mated with wild-type counterparts, and the resulting G1 larvae were screened for green fluorescence using the NIGHTSEA system (EMS, USA). Positive lines were maintained as heterozygous through outcrosses with wild-type (HSDdsx) or as homozygotes (*vasa*-Cas9) via intercrosses.

### Crosses and phenotypes

To evaluate drive efficiency, heterozygous transgenic mosquitoes were crossed with their siblings or wild-type mosquitoes. Progeny were screened for fluorescence at the larval stage. Drive carriers (displaying fluorescence) and non-drive individuals (lacking fluorescence) were segregated and reared separately until adulthood. Subsequently, individuals were sexed using a stereo microscope (Olympus, USA).

In investigating the potential enhancement of HSDdsx drive efficiency with an additional *vasa*-Cas9 source, HSDdsx and *vasa*-Cas9 adults were crossed to generate double-heterozygous lines containing both drive alleles. The resulting double-heterozygous males were then crossed with wild-type females, and their offspring were phenotyped.

### Genotyping

Genomic DNA was extracted using either DNAzol Reagent (Invitrogen, UK) or the Animal Genomic DNA Quick Extraction Kit (Beyotime, China). PCR reactions were performed with Q5 High-Fidelity DNA Polymerase (NEB, UK). Genomic integration of both constructs were confirmed through PCR and Sanger sequencing. For genotyping of each target gene, a primer pair covering the target region was designed to detect possible end joining-induced resistant alleles. Primers (708: 5’-atcttgctcctcacttgccc −3’; 710: 5’-ggtgtcgcccactccttaaac −3’) were designed for amplifying a 539 bp region covering *dsx* target sites, and primers (711: 5’-tcaaagctgccacggatctc −3’; 714: 5’-aacccagactatgtgaaggatg −3’) were used to amplify a 655 bp fragment of *GUY1* specifically present in Y chromosome. Another pair of primers (565: 5’-tcgtatcaacaactgtctgaacgagctg-3’; 39: 5’-tgaaggatggccggccaatc-3’) were designed to amplify 557 bp covering a fragment of *hairy* genomic sequence.

### Amplicon sequencing

Heterozygous HSDdsx adults were crossed with each other, and their offspring were subsequently phenotyped. A total of 100 adults, encompassing both individuals with and without the drive allele were pooled for genomic DNA extraction. The target region was amplified using primers 581 (5’-agaagatgaggctcttgatcttgatc-3’) and 582 (5’-agaactatcgaagaattcggttcacc-3’). Subsequently, PCR products were purified using the Zymoclean Gel DNA Recovery Kit (Zymo Research, USA) to prepare for deep sequencing conducted by Genwiz. Sequencing data has been uploaded into Github (https://github.com/jchamper/ChamperLab/tree/main/Anopheles-stephensi-dsx-Drive).

### Modeling

A previously developed mosquito-specific SLiM model^32–35^, which simulated various life stages of mosquitoes, was applied to analyze population dynamics by releasing different combinations of HSDdsx into a wild population. Simulations were run for 317 weeks post release, with 3.167 weeks representing one generation. The wild-type population was allowed to equilibrate for 10 generations before releasing drive carriers.

For HSDdsx, we set up a one-time release of male heterozygotes with various combinations of low-density growth rate and female heterozygote fitness. The low-density growth rate represents female fecundity under optimal conditions without competition, while the female fertility reflects the fitness of female drive heterozygotes. Other default parameters were set up based on our experimental results. The data of its simulated drive allele frequency, r2 resistance allele frequency, total fertile female number, and total adult number in each week were collected.

We also conducted a combined drive continuous release, assuming a fitness-neutral *vasa*-Cas9 allele and a HSDdsx drive lacking Cas9. In this simulation, mosquitoes that were heterozygous for the split suppression drive allele and homozygous for the Cas9 allele were continuously released into the population every week. If the population was not eliminated, we recorded the number of fertile females and drive female frequency. If population elimination took place, we recorded the time needed for successful population suppression.

Considering that both drives used a gRNA multiplexing strategy to prevent the generation of functional r1 alleles, only non-functional r2 alleles were modeled. Each simulation was independently ran five times, and data was collected for figure generation with Python. The default parameters for both drives are listed in Table S1, and corresponding SLiM scripts and data can be found in Github (https://github.com/jchamper/ChamperLab/tree/main/Anopheles-stephensi-dsx-Drive).

## Results

### Construction of a homing suppression drive HSDdsx

We built a suppression drive (HSDdsx) in *A. stephensi*, specifically disrupting the female exon of the haplosufficient gene *dsx* (Figure 1A). It was expected that the female offspring inheriting two disrupted alleles would be sterile and eliminated from the population (Figure 1B). Our suppression drive design was inspired by a previous study in *A. gambiae*^24^ but incorporated several enhancements. Firstly, we employed two gRNAs, targeting closely located sequences around the boundary, each regulated by distinct Pol III promoters (U6a and 7SK) and positioned in opposite directions to prevent recombination and deletion of one of the genes. The proximity of these gRNA target sites was intentional to further reduce functional resistance and potential homology-direct repair efficiency reduction. Second, Cas9 expression was driven by a *nanos* promoter, which may reduce fitness costs^32^ caused by somatic disruption of *dsx* in heterozygous females. Following the injection of 1110 eggs, 58 adults survived, and upon crossing with wildtypes, only one EGFP-positive male was recovered. Subsequently, this male was crossed with wild-type females to establish the HSDdsx transgenic line. Morphological analysis revealed no observable differences between heterozygous drive carriers and wild-type mosquitoes.

**Figure 1.**
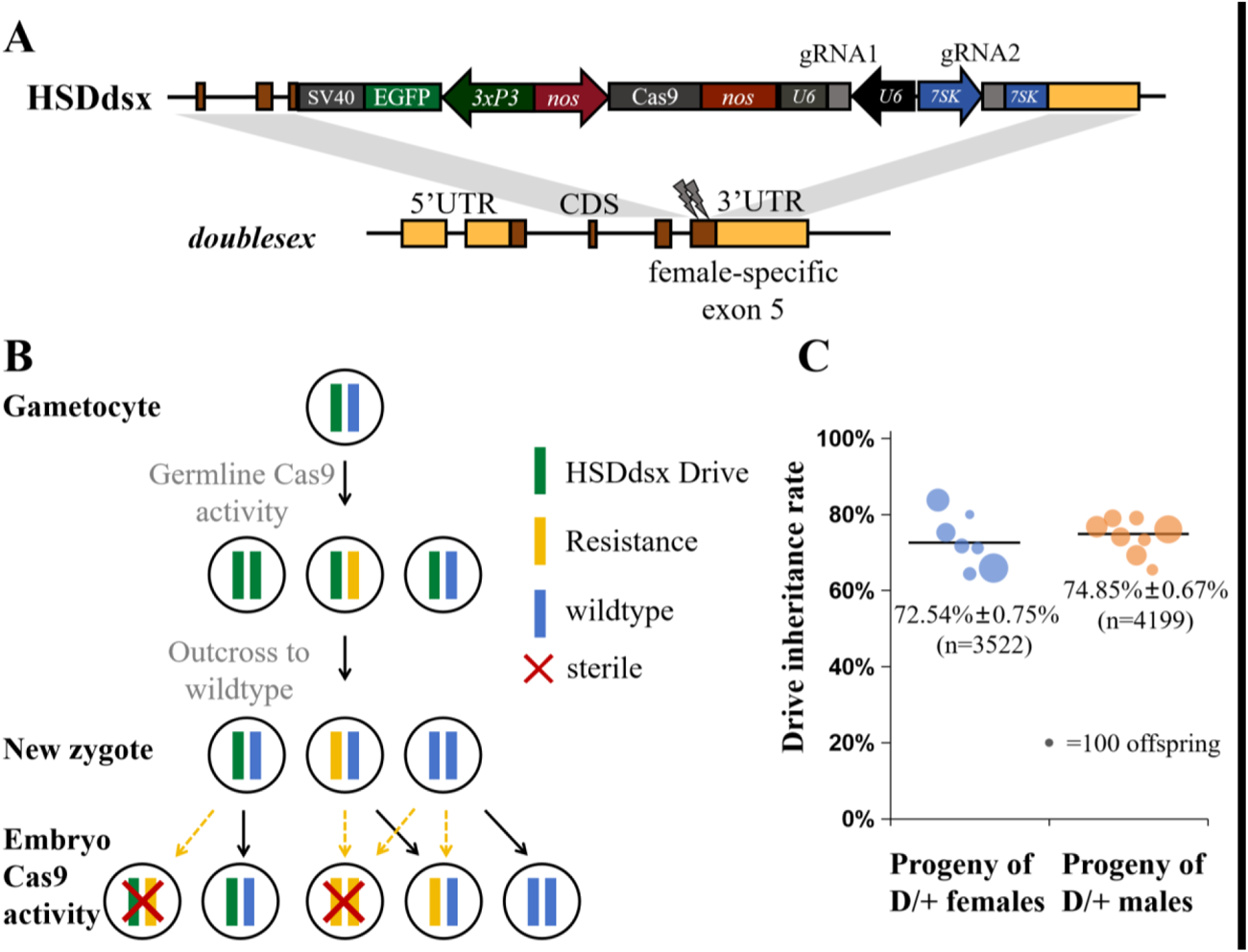
Design and performance of the suppression drive HSDdsx. (**A**) HSDdsx is inserted into the female-specific intron 4-exon 5 boundary of *dsx*. The drive element contains a *nanos-Cas9* cassette, a 3xP3-EGFP-SV40 marker, and two gRNAs under the control of U6a and 7SK promoters. (**B**) A drive allele will express Cas9 to cut the wild-type allele in the germline, after which the wild-type allele is either converted to drive allele, or disrupted via end-joining. Maternally deposited Cas9/gRNA can cut wild-type alleles in the embryo, followed by disruption through end-joining. Females carrying two non-functional alleles (drive or resistance) will be sterile. (**C**) Drive heterozygotes were outcrossed with wild-type, and their offspring were phenotyped for EGFP fluorescence, indicating the rate of drive inheritance. The size of the dots indicates the relative sample size of larval progeny from a single cross batch. “n” indicates the total number of offspring in each group. The mean and standard error of the mean (SEM) are displayed.

### Drive efficiency test of HSDdsx

To assess the drive efficiency of HSDdsx, we initiated crosses between drive heterozygotes (G0) and wildtypes. The generated heterozygous drive carriers (G1) were then either outcrossed with wildtypes or intercrossed with heterozygous siblings. Their G2 larvae progeny were screened for fluorescence, and morphological phenotyping was conducted upon their emergence as adults. The drive inheritance rate was measured as the percentage of EGFP-expressing larvae. Our findings revealed slightly higher drive inheritance rates for HSDdsx when G1 drive parents were males (74.85%±0.67%) compared to females (72.54%±0.75%, *p*=0.022, Fisher’s Exact Test). These correspond to drive conversion rates of 49.70%±1.34% and 45.09%±1.50%, respectively. Notably, both drive male and female groups exhibited drive inheritance rates significantly higher than the 50% Mendelian expectation (*p*<0.0001, Binomial Test). For the crosses between male and female heterozygotes, the drive carrier rate among offspring was 94.60%±0.43%, consistent with the inheritance rates from each heterozygote parent (Figure 1C, Data Set S1). This result suggested the expression of Cas9 driven by *nanos* promoter and two gRNAs respectively driven by U6 and 7SK promoters were functional in causing biased inheritance of the drive allele, though the drive efficiency was lower than a similar suppression drive in the closely related species *Anopheles gambiae*^24^.

Fertility tests for drive females, originating from either paternal or maternal drive, showed a significantly lower hatch rate when the G0 drive carrier was female (56.54%±3.59%, *p*=0.0012, Fisher’s Exact Test), while no significant difference was observed between G0 drive males (74.08%±2.24%) and wild-type controls (85.75%±1.87%) (*p*=0.0698, Fisher’s Exact Test). This suggests a potential fitness cost associated with maternal drive, though the fertility test could also be influenced by population density or batch effects (Data Set S2).

Phenotyping of offspring resulting from the outcross of heterozygotes and wildtypes revealed normal sex morphological features, of which male had plumose antennae and downward facing claspers in its external genitalia while female had pilose antennae and cerci (Figure 2). We randomly collected 15 offspring for genotyping, which showed no resistance alleles in either drive or non-drive individuals. However, when drive heterozygotes were intercrossed, 60% of drive-carrying offspring showed three various intersex phenotypes, which we termed as intersex XX, intersex XY, and intersex-90 XY. Intersex XX was genetically female, showing less-bushy antennae and upward rotated claspers. Intersex XY and intersex-90 XY were both genetically males and had male-like bushy antennae, but the former had upward claspers while the later exhibited claspers that were twisted 90 degrees (Figure 2 and S1). Genotyping revealed that all of the intersex XX and intersex-90 XY mosquitoes were drive homozygous females and males, respectively, and most intersex XY mosquitoes were also homozygous males. Note that two out of 14 intersex XY mosquitoes had mosaic wildtype/resistance alleles, with the wild-type sequence dominant (in addition to one drive allele). While it is unlikely that resistance alleles could disrupt male expression, this may be possible. More likely, these mosquitoes were misidentified because wild-type males also exhibit up-toward claspers shortly after emergence, but they eventually rotate 180 degrees within ∼48 hours post-eclosion. The upward claspers of these two individuals might have eventually been able to rotate in an extended time window (though phenotyping was conducted five days post-eclosion), or they may have had a developmental defect.

**Figure 2.**
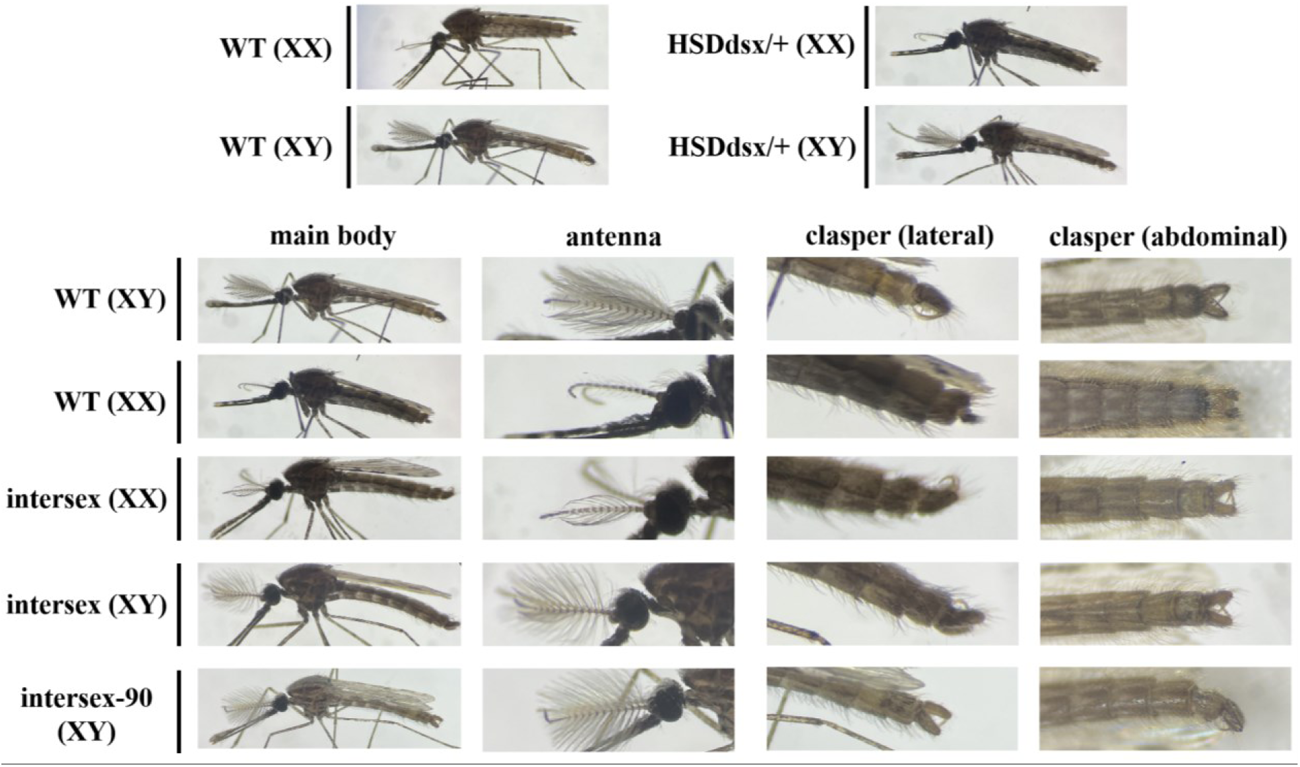
The morphology analysis of HSDdsx line. Heterozygous drive males and females show identical normal morphology with wildtypes, while various intersex phenotypes were observed in null mutants. XX and XY indicate genetic females and males, respectively, and all the intersex mosquitoes were drive homozygotes, as confirmed by genotyping.

These male intersex mosquitoes were respectively pool-crossed to wild-type females, each with 5 males and 15-20 females and three cages each for intersex and intersex-90 males, but none of them produced any progeny. Our findings suggest that *dsx* is a haplosufficient gene in *A. stephensi* and that insertion of the drive construct into the female-specific intron 4-exon 5 boundary disrupted *dsx* expression and introduced sterility in both males and females.

### Potential resistance in HSDdsx

To confirm if resistance emerged in our suppression drive line, deep sequencing was conducted around the closely spaced gRNA target sites. 100 drive and non-drive offspring from drive heterozygous parents were pooled for deep sequencing. The result showed a very low level of cleavage at both gRNA target sites, with only 0.6568% and 1.3813% reads modified in gRNA1 and gRNA2 targets, respectively. Probably only 1-2 mosquitoes had a resistance allele (with a cut at the second gRNA under control of the 7SK promoter) that was formed from the germline or at the zygote stage (and thus inherited by all cells in the offspring). Most detected resistance alleles were found in a small proportion and were thus likely formed by somatic expression or mosaic cleavage from maternal Cas9. None of these mutations seemed to be in-frame, making it unlikely that any would be functional (Figure S2). As expected from the low cut rates, the frequency of simultaneous cut of both gRNAs was even lower (<0.00021%). This result reveals that both gRNAs were active and that resistance rates were overall very low, preventing a thorough assessment of functional versus nonfunctional resistance.

### Modeling of HSDdsx

To assess HSDdsx performance at population level, we applied a SliM mosquito model to analyze allele frequency and population size on a weekly basis following a 5% introduction of heterozygous drive males into the population. We compared our drive performance in two scenarios where male homozygotes are fertile and sterile, respectively. In addition, two different low-density growth rates to simulate different potential population dynamics.

Our results demonstrated a consistent increase in drive allele frequency until reaching equilibrium (Figure 3A). However, while all drives affected the population size, only the population with fertile male homozygotes and lower low-density growth rate was successful at eliminating the target population (Figure 3B). The drive’s genetic load (suppressive power) was too weak to overcome a low-density density growth rate of 6, and sterile homozygous males substantially reduced suppression power. This was because nonfunctional resistance alleles gained an advantage in this scenario over the drive allele (only drive homozygous males are sterile, while fort females, any combination of drive and nonfunctional resistance alleles are sterile). Notably, the population size expanded when low-density growth rate was high (Figure 3B). This was because the reduction in larvae competition (there were fewer eggs because of sterile females and reduced fertility females) allowed more larvae to survive.

**Figure 3.**
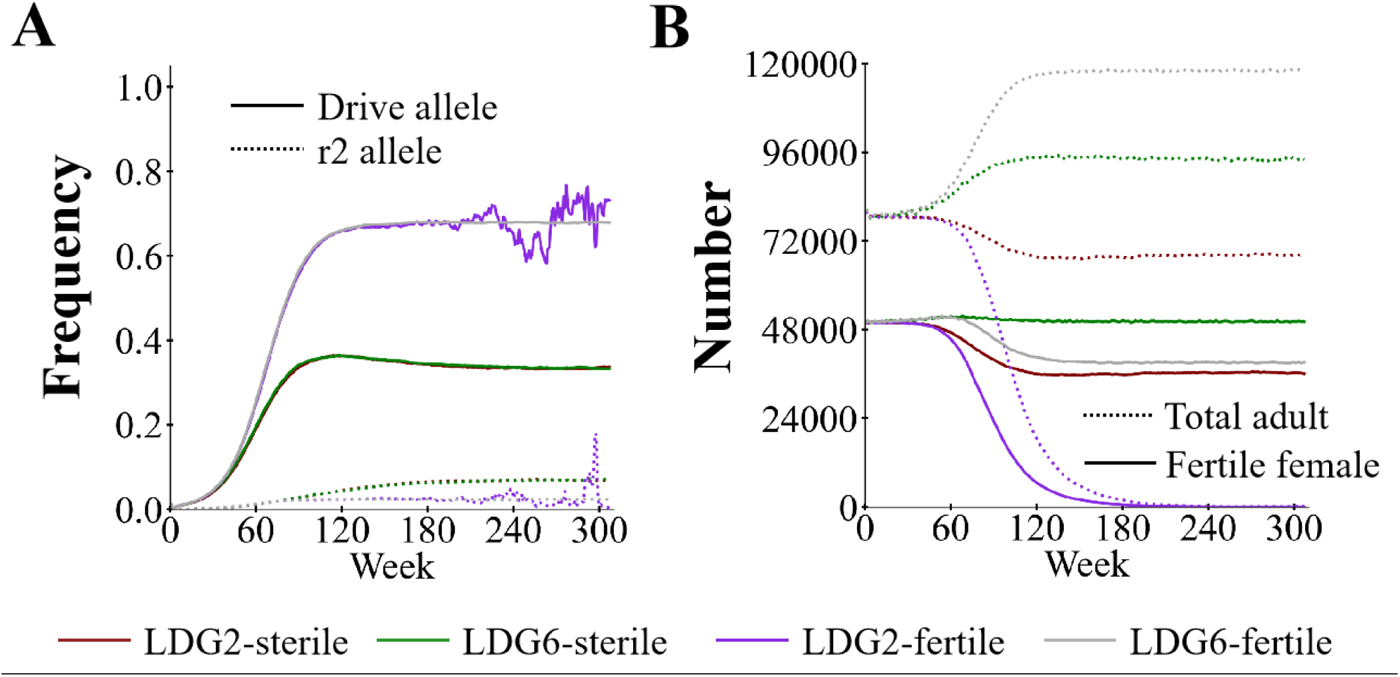
HSDdsx drive performance in a mosquito model. Drive heterozygous males were released into the population at 5% frequency. Drive performance parameter were based on experimental results (Table S1), with and without homozygous male sterility, as well as low and high low-density growth rate (LDG). (**A**) Allele frequencies and (**B**) average number of fertile females are displayed. Each simulation was repeated five times, and average values are shown in the figure. r2 indicates nonfunctional resistance alleles.

### Combination of HSDdsx and *vasa*-Cas9

In addition to HSDdsx, we generated another *A. stephensi* transgenic line, which was based on a “toxin-antidote” modification drive in a previous *D. melanogaster* study^13^. However, this line was not able to bias its inheritance, likely due to low cleavage efficiency at its *hairy* target site. We thus employed it as a *vasa*-Cas9 line. To further investigate the performance of both lines, we first crossed HSDdsx with *vasa*-Cas9 (G0) to generate double heterozygous individuals (G1). Though both lines had the 3xP3-EGFP marker, their fluorescence patterns were different, enabling the reliable identification of each allele, even when both were present in the same individual (Figure 4B). The inheritance rates of HSDdsx and *vasa*-Cas9 in G1 were 79.77%±1.44% and 50.58%±1.79% when HSDdsx females were crossed with *vasa*-Cas9 males. This was consistent with the results of drive efficiency tests of each line. *Vasa* promoter has been known active in both somatic tissues and maternal deposition^36^. To avoid Cas9 deposition from maternal *vasa*-Cas9 allele, only the double heterozygotes from HSDdsx female and *vasa*-Cas9 male cross were selected to cross with wildtypes for further drive efficiency test. Notably, all the double heterozygotes were confirmed as either intersex or morphological males, without any females being identified, indicating complete masculinization in drive females due to somatic cutting of *dsx*. Additionally, we genotyped seven of these double heterozygous males and found mosaic mutations in both the *dsx* gRNA target sites of six, indicating high somatic expression of Cas9.

Double heterozygous males were further crossed to wildtypes to create G2 offspring. The phenotyping of G2 offspring revealed a significant boost in the HSDdsx drive inheritance to 100%, while the inheritance of *vasa*-Cas9 was 42.49%±1.78% (Data Set S4). This showed that an extra Cas9 source was sufficient to induce higher germline cleavage and conversion of HSDdsx. The *vasa*-Cas9 inheritance was slightly reduced (even compared to previous crosses of *vasa*-Cas9 and wild-type), which could be potentially explained by the high fitness cost of some individuals carrying both drive alleles, leading to death in early stages, or perhaps *vasa*-Cas9 carrying sperm were less competitive than wild-type sperms.

We observed that females with one HSDdsx and one *vasa*-Cas9 allele exhibited intersex phenotypes, but the extent of intersex characteristics varied. Females with *vasa*-Cas9 fathers and HSDdsx mothers displayed up-toward claspers (as shown in intersex XX), whereas female progeny of males heterozygous for both HSDdsx and *vasa*-Cas9 showed more pronounced abnormalities in external genitalia (Figure S3A). Genotyping of these latter individuals carrying both HSDdsx and *vasa*-Cas9 alleles confirmed their status as females heterozygous for HSDdsx and mutated alleles at the *dsx* targets (Figure S3B).

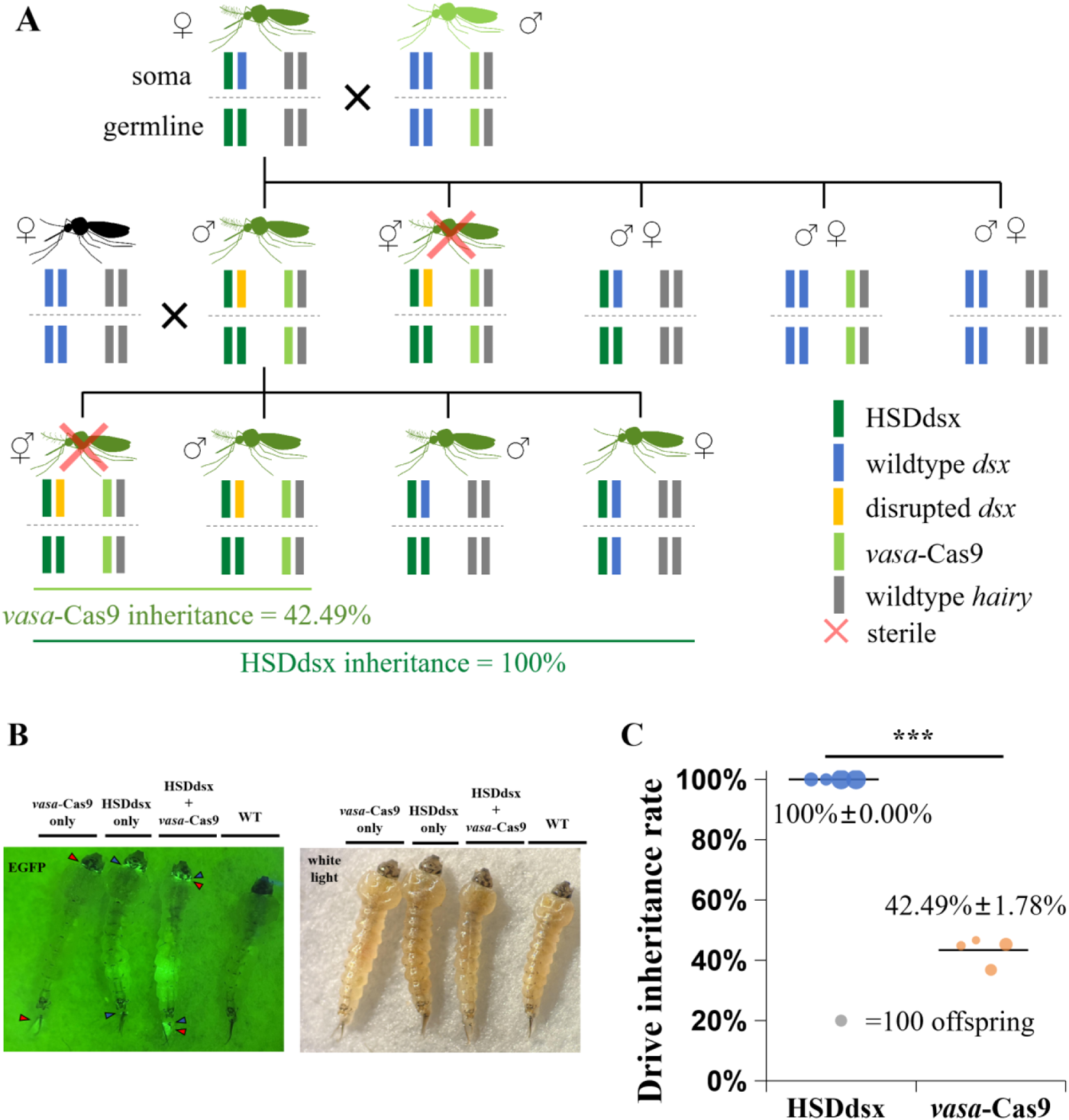
scheme of HSDdsx and *vasa*-Cas9 lines and the drive inheritance in offspring. (**A**) HSDdsx females and *vasa*-Cas9 males were crossed to generate double het males, which were crossed to wild-type for assessing drive inheritance in offspring. Disrupted dsx can come from drive conversion and/or resistance allele formation in somatic cells. (**B**) The left panel picture was taken with an EGFP filter, and the right panel was under white light. Triangles with different colors were used to present the differences of fluorescence patterns among drive lines. Wild-type (WT) mosquitoes are also displayed for comparison. **(C)** In the offspring of double heterozygous drive males and wild-type females, HSDdsx inheritance was boosted to 100%, while *vasa*-Cas9 inheritance was 42.49%, significantly lower than HSDdsx (*p*<0.0001***, Fisher’s exact test).

### Modeling of the combination of HSDdsx and *vasa*-Cas9 for SIT-like population suppression

Based on the results of the combination crosses, we propose a SIT-like strategy for population suppression and assessed its capacity via our SLiM mosquito modeling. This combination drive consists of two components located in unlinked loci, a gRNA-only homing allele targeting *dsx* and an unlinked Cas9 allele. This allows the system to be self-limiting, though use of our actual lines would allow slightly higher suppressive power at the cost of potential spread of the weak but unconfined HSDdsx into nontarget populations. Our modeling result showed that the population could be successfully suppressed when the release ratios were higher than 0.8 (Figure 5A). When release ratio was 0.8, 20% of the simulated populations could be successfully suppressed. The drive female frequency and the number of fertile females eventually reached equilibrium when release ratios were lower (Figure 5B and C). Having a higher release ratio sped up suppression, but this had rapidly decreasing returns (Figure 5D). These results demonstrated that our combined drive can be applied as an efficient self-limiting suppression strategy, with substantially higher efficiency than SIT and RIDL strategies^33^.

**Figure 4.**
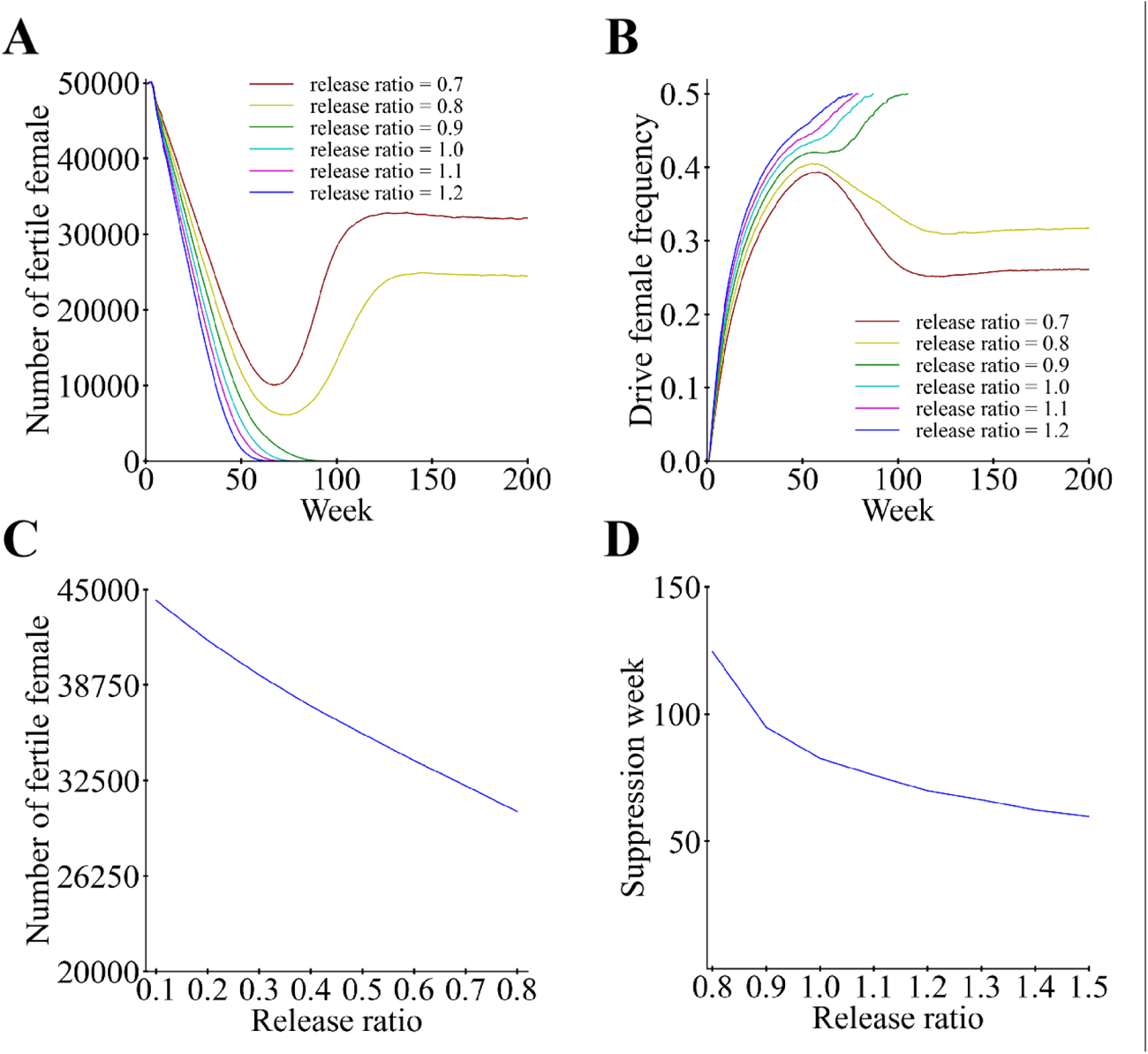
Combination drive model. Mosquitoes carrying heterozygous drive allele and homozygous Cas9 allele were released into the population every week. (**A**) and (**B**) show the number of fertile females and the drive female frequency, respectively, over time at different release ratios. (**C**) Number of fertile females at various release ratios when the population is not eliminated. (**D**) Actual time needed to suppress the population when elimination is possible. Note that for the release ratio of 0.8, population elimination occurred in four out of 20 simulations, so this release level is shown in C and D to assess both situations.

## Discussion

In this study, we developed the first homing suppression gene drive (HSDdsx) in *A. stephensi*. Our experimental findings support several conclusions: 1) the *nanos-Cas9* in HSDdsx was functional but not strong enough to induce high germline cut rate and conversion rate; 2) the multiplexed gRNA expression driven by U6 and 7SK promoters was effective, and modeling indicates that this is likely to substantially reduce functional resistance allele formation rates^37^; 3) resistance allele formation from germline-restricted *nanos-Cas9* was minimal, though this could have been due to low expression in general; 4) the expression of *vasa*-Cas9 in the transgenic line was high, supporting 100% drive conversion efficiency in HDSdsx.

Our suppression drive was inspired by the previous study in *A. gambiae*, with improvements including gRNA multiplexing, the use of a different germline promoter *nanos*, and an insect codon-optimized Cas9 sequence. Suppression drives are generally more sensitive to fitness costs caused by leaky somatic Cas9 expression or resistance allele formation from early embryonic activity of Cas9/gRNA, as reported in *D. melanogaster*^28,30^ and *Anopheles* mosquitoes^27,38^. Even though *A. stephensi* and *A. gambiae* are relatively closely related, the *A. stephensi nos* promoter was substantially less effective in *A. stephensi* than the *A. gambiae nos* promoter in *A. gambiae*, though one caveat to this is that the *A. gambiae* examples were at a different target site^26,38^. This difference could be attributed to lower expression of Cas9, which may have been due to the limited size of our *nos* promoter, which could have been missing regulatory sequences. *nos* may also simply have lower expression in *A. stephensi* than *A. gambiae*. Regulatory elements flanking the drive insertion site could also have affected expression. To enhance Cas9 germline expression and genetic load of the drive, future experiments can explore the activity of *nos* promoters in different lengths, target different genes or loci, or test other germline promoters (e.g. the *zpg* promoter, which also has excellent performance in *A. gambiae*^24,38^).

Based on publications in *A. gambiae*, disrupting the female-specific exon of *dsx* converted only homozygous null-mutant females (either with or without drive) into an intersex phenotype causing sterility, while males remained healthy and fertile^24^. Our observations in closely related *A. stephensi* showed that homozygous males also showed intersex phenotypes (Figure 2). The existence of intersex homozygous males can be potentially explained by the insertion of the drive construct, affecting the transcription level and potentially the splicing of male-specific *dsx* exons, which are downstream of the gene drive. This effect on fitness may not be obvious in heterozygous males because the wild-type allele is haplosufficient. It remains unclear why this occurred in our *A. stephensi* drive but not in *A. gambiae* studies. Perhaps *nanos* regulatory elements terminated transcription at higher rates than *zpg* elements, despite being in the opposite orientation of *dsx*. An similar intersex phenotype in homozygous males has also been observed in *Drosophila suzukii*^22^, while other reports in *D. suzukii* and *D. melanogaster* also showed the dominant sterility in drive females, illustrating the complexity of sex-specific *dsx* expression^22,39^. These, together with the current study, demonstrate the necessity of more detailed assessments of both male and female in such suppression drives targeting *dsx*. Homozygous male sterility could substantially reduce overall drive efficiency and make the drive more vulnerable to chasing^40^.

Resistance alleles could be classified into functional (r1) and non-functional (r2) alleles. While germline-restricted promoters can help in reducing total resistant allele formation, gRNA multiplexing is a useful method to reduce the fraction of r1 alleles, which is essential for suppression drive success. The separate gRNA expressing cassettes (with different promoters, U6 and 7SK) used in HSDdsx demonstrated strong gRNA expression capacity, providing an alternative tool for future gRNA multiplexing. While increased numbers of gRNAs can eventually reduce drive frequency^37^, it is more important to minimize functional resistance. In this study, the reduced drive efficiency was not likely due to use of a second gRNA because the total resistance allele formation rate was very low, and failed drive conversion due to gRNA multiplexed is likely to produce resistance alleles^37^. Further, if resistance alleles produced by the multiple gRNAs turn out to be dominant sterile, as seen in *dsx* for *D. melanogaster*^39^ and for at least some alleles in *A. gambiae*^41^, then the drive may prove to have better performance than a standard homing suppression drive, even with male homozygous sterility.

Our research yields insights for the advancement of efficient and environmentally friendly pest control tools aimed for disrupting disease transmission. It suggests that constructing high-efficiency drives in *A. stephensi* could be achieved through modest modifications to our existing constructs. For the homing suppression drive, a stronger germline promoter can be used to drive Cas9 expression, which could increase drive efficiency and potentially suppress the population even with homozygous sterility in both sexes. It may also be possible to restore male homozygote fertility.

## Acknowledgements

This study was supported by grants from the National Science Foundation of China (32302455 and 32270672). We also thank Peking University and the Center for Life Sciences for providing laboratory startup support.

## Supplemental Information

**Table S1.** Default parameters for the mosquito model.

| Drive | Name | Default value |
| --- | --- | --- |
| HSDdsx | Number of adult females | 50000 |
|  | Drop rate (reflecting release size) | 0.05 |
|  | Female drive conversion rate | 0.4509 |
|  | Male drive conversion rate | 0.497 |
|  | Female drive embryo resistance cut rate | 0.01 |
|  | Male drive embryo resistance cut rate | 0 |
|  | Female germline resistance allele formation rate | 0 |
|  | Male germline resistance allele formation rate | 0 |
|  | Female heterozygote fitness | 0.796 |
|  | Male only release | True |
|  | Heterozygote release | True |
| Combination | Number of adult females | 50000 |
|  | Drive conversion rate | 0.99 |
|  | Germline resistance allele formation rate | 0 |
|  | Embryo resistance cut rate | 0 |

**Figure S1.**
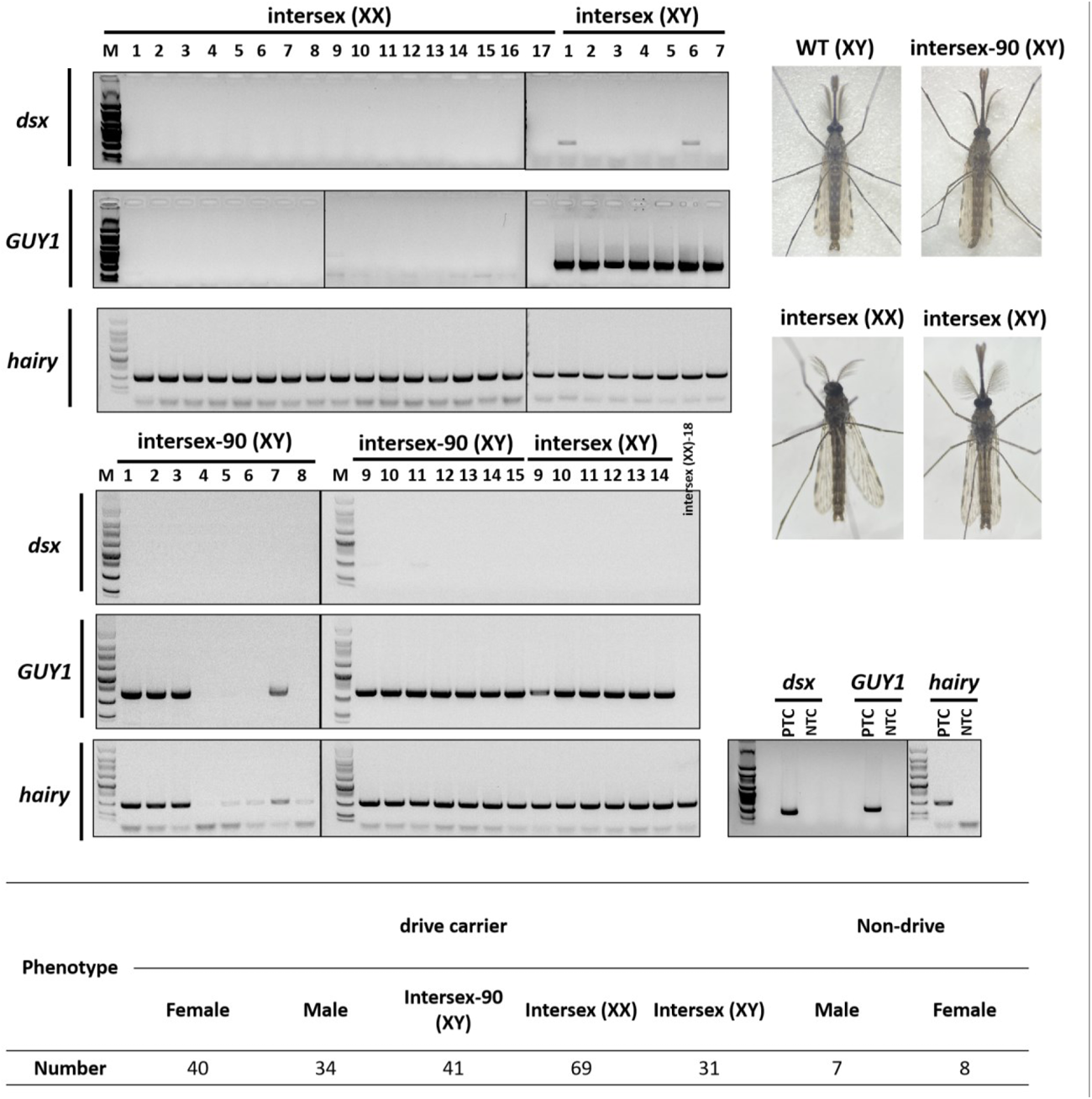
Genotyping of intersex individuals. Drive males and drive females were crossed, and their offspring with EGFP were collected for phenotyping as well as genotyping. Pairs of primers were designed to amplify the *dsx* target region (539 bp) and a Y chromosome gene *GUY1* (655 bp) for determining the genotype and genetic sex of samples, while another primer pair was used to amplify a *hairy* fragment (557 bp) as the positive control for indicating genomic DNA template quality. Three different intersex phenotypes were observed among drive-carrying mosquitoes. Intersex (XX) and intersex (XY) were genetically female and male, respectively. Intersex-90 (XY) was genetically male, which had normal antenna but a twist of 90 degrees in the male genitalia in the abdomen, showing an intermediate phenotype between intersex and wild-type. The absence of *GUY1* PCR product in intersex-90 samples 4, 5, 6 and 8 was likely due to poor DNA quality (indicated by much weaker *hairy* bands). PCT: positive control with wild-type gDNA template. NCT: negative control with water template.

**Figure S2.**
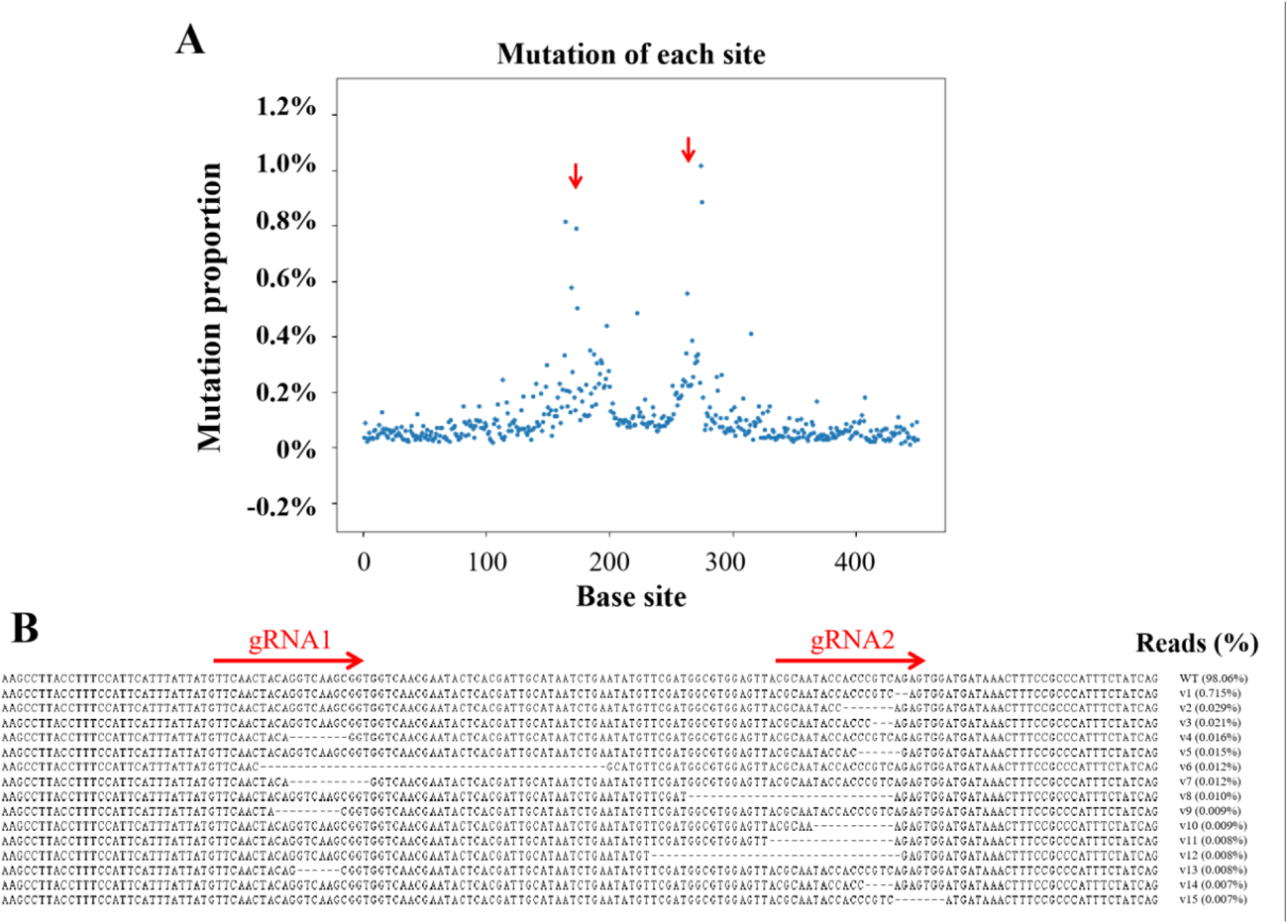
Deep sequencing reveals mutations at gRNA target sites. (**A**) The fraction of mutations at each nucleotide differing from wildtype. (**B**) Examples of reads obtained in deep sequencing and their corresponding frequency. Red arrows indicate gRNA locations.

**Figure S3.**
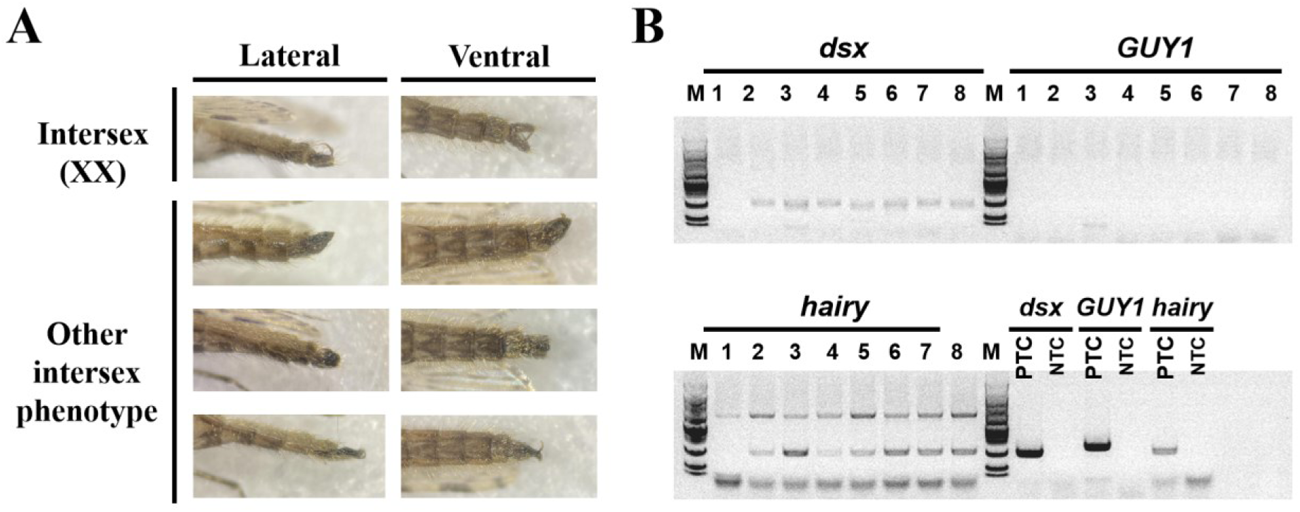
The abnormalities observed in HSDdsx+*vasa*-Cas9 females. (**A**) Pictures of external genitalia of intersex individuals. (**B**) Genotyping of mosquitoes with other intersex phenotypes. Three pairs of primers were used to identify presence of a wild-type or resistance allele at *dsx*, the presence of a Y chromosome (based on the *GUY1* gene), and the quality of the genomic templates (based on the *hairy* gene). The amplicons of *dsx* were cleaned-up for sequencing, showing mutations in target sites. PTC means positive control using wild-type DNA template, and NTC means negative control with water as template. Upper bands in *hairy* were non-specific products. The genotyped abnormal individuals were all heterozygous drive females containing mosaic *dsx* mutations.

## References

1. Reid, M. C. & McKenzie, F. E. The contribution of agricultural insecticide use to increasing insecticide resistance in African malaria vectors. Malar J 15, 107 (2016).

2. Robinson, A. S. Genetic Basis of the Sterile Insect Technique. in Sterile Insect Technique: Principles and Practice in Area-Wide Integrated Pest Management (eds. Dyck, V. A., Hendrichs, J. & Robinson, A. S.) 95–114 (Springer Netherlands, Dordrecht, 2005). doi:10.1007/1-4020-4051-2_4.

3. Black, W. C., Alphey, L. & James, A. A. Why RIDL is not SIT. Trends Parasitol 27, 362–370 (2011).

4. Li, M. et al. Suppressing mosquito populations with precision guided sterile males. Nat Commun 12, 5374 (2021).

5. Fu, G. et al. Female-specific flightless phenotype for mosquito control. Proc Natl Acad Sci U S A 107, 4550–4554 (2010).

6. Alphey, L. Genetic Control of Mosquitoes. Annual Review of Entomology 59, 205–224 (2014).

7. Marinotti, O. et al. Development of a population suppression strain of the human malaria vector mosquito, *Anopheles stephensi*. Malaria Journal 12, 142 (2013).

8. Adams, K. L., et al. *Wolbachia cifB* induces cytoplasmic incompatibility in the malaria mosquito vector. Nat Microbiol 6, 1575–1582 (2021).

9. McNamara, C. J. et al. Transgenic expression of cif genes from Wolbachia strain wAlbB recapitulates cytoplasmic incompatibility in *Aedes aegypti*. Nat Commun 15, 869 (2024).

10. Zheng, X. et al. Incompatible and sterile insect techniques combined eliminate mosquitoes. Nature 572, 56–61 (2019).

11. Burt, A. Site-specific selfish genes as tools for the control and genetic engineering of natural populations. Proc Biol Sci 270, 921–928 (2003).

12. Champer, J. et al. A CRISPR homing gene drive targeting a haplolethal gene removes resistance alleles and successfully spreads through a cage population. Proceedings of the National Academy of Sciences 117, 24377–24383 (2020).

13. Champer, J. et al. A toxin-antidote CRISPR gene drive system for regional population modification. Nat Commun 11, 1082 (2020).

14. Faber, N. R. et al. Improving the suppressive power of homing gene drive by co-targeting a distant-site female fertility gene. 2023.12.07.570117 Preprint at 10.1101/2023.12.07.570117 (2023).

15. Pfitzner, C. et al. Progress Toward Zygotic and Germline Gene Drives in Mice. The CRISPR Journal 3, 388–397 (2020).

16. Grunwald, H. A. et al. Super-Mendelian inheritance mediated by CRISPR-Cas9 in the female mouse germline. Nature 566, 105–109 (2019).

17. Oberhofer, G., Johnson, M. L., Ivy, T. & Hay, B. A. Cleave and Rescue gamete killers create conditions for gene drive in plants. bioRxiv 2023.10.13.562303 (2023) doi:10.1101/2023.10.13.562303.

18. Liu, Y., Jiao, B., Champer, J. & Qian, W. Overriding Mendelian inheritance in Arabidopsis with a CRISPR toxin-antidote gene drive that impairs pollen germination. 2023.10.10.561637 Preprint at 10.1101/2023.10.10.561637 (2023).

19. DiCarlo, J. E., Chavez, A., Dietz, S. L., Esvelt, K. M. & Church, G. M. Safeguarding CRISPR-Cas9 gene drives in yeast. Nat Biotechnol 33, 1250–1255 (2015).

20. Walter, M. et al. Viral gene drive spread during herpes simplex virus 1 infection in mice. 2023.12.07.570711 Preprint at 10.1101/2023.12.07.570711 (2024).

21. Xu, X. et al. Toward a CRISPR-Cas9-Based Gene Drive in the Diamondback Moth *Plutella xylostella*. CRISPR J 5, 224–236 (2022).

22. Yadav, A. K. et al. CRISPR/Cas9-based split homing gene drive targeting *doublesex* for population suppression of the global fruit pest *Drosophila suzukii*. Proc Natl Acad Sci U S A 120, e2301525120 (2023).

23. Anderson, M. A. E. et al. A multiplexed, confinable CRISPR/Cas9 gene drive can propagate in caged *Aedes aegypti* populations. Nat Commun 15, 729 (2024).

24. Kyrou, K. et al. A CRISPR-Cas9 gene drive targeting *doublesex* causes complete population suppression in caged *Anopheles gambiae* mosquitoes. Nat Biotechnol 36, 1062–1066 (2018).

25. Adolfi, A. et al. Efficient population modification gene-drive rescue system in the malaria mosquito *Anopheles stephensi*. Nat Commun 11, 5553 (2020).

26. Carballar-Lejarazú, R. et al. Dual effector population modification gene-drive strains of the African malaria mosquitoes, *Anopheles gambiae* and *Anopheles coluzzii*. Proceedings of the National Academy of Sciences 120, e2221118120 (2023).

27. Hammond, A. et al. A CRISPR-Cas9 gene drive system targeting female reproduction in the malaria mosquito vector *Anopheles gambiae*. Nat Biotechnol 34, 78–83 (2016).

28. Du, J. et al. New germline Cas9 promoters show improved performance for homing gene drive. 2023.07.16.549205 Preprint at 10.1101/2023.07.16.549205 (2023).

29. Oberhofer, G., Ivy, T. & Hay, B. A. Cleave and Rescue, a novel selfish genetic element and general strategy for gene drive. Proc Natl Acad Sci U S A 116, 6250–6259 (2019).

30. Yang, E. et al. A homing suppression gene drive with multiplexed gRNAs maintains high drive conversion efficiency and avoids functional resistance alleles. G3 (Bethesda) 12, jkac081 (2022).

31. Simoni, A. et al. A male-biased sex-distorter gene drive for the human malaria vector *Anopheles gambiae*. Nat Biotechnol 38, 1054–1060 (2020).

32. Champer, S. E., Kim, I. K., Clark, A. G., Messer, P. W. & Champer, J. *Anopheles* homing suppression drive candidates exhibit unexpected performance differences in simulations with spatial structure. Elife 11, e79121 (2022).

33. Zhu, Jinyu, Chen, Jingheng, Liu, Yiran, Xu, Xuejiao, & Champer, Jackson. Population suppression with dominant female-lethal alleles is boosted by homing gene drive. bioRxiv 2023.12.05.570109 (2023) doi:10.1101/2023.12.05.570109.

34. Liu, Y., Teo, W., Yang, H. & Champer, J. Adversarial interspecies relationships facilitate population suppression by gene drive in spatially explicit models. Ecol Lett 26, 1174–1185 (2023).

35. Haller, B. C. & Messer, P. W. SLiM 4: Multispecies Eco-Evolutionary Modeling. Am Nat 201, E127–E139 (2023).

36. Gantz, V. M. et al. Highly efficient Cas9-mediated gene drive for population modification of the malaria vector mosquito *Anopheles stephensi*. Proc Natl Acad Sci U S A 112, E6736–E6743 (2015).

37. Champer, S. E. et al. Computational and experimental performance of CRISPR homing gene drive strategies with multiplexed gRNAs. Science Advances 6, eaaz0525 (2020).

38. Hammond, A. et al. Regulating the expression of gene drives is key to increasing their invasive potential and the mitigation of resistance. PLoS Genet 17, e1009321 (2021).

39. Weizhe Chen, Jialiang Guo, Yiran Liu, & Jackson Champer. Population suppression by release of insects carrying a dominant sterile homing gene drive targeting *doublesex* in *Drosophila*. bioRxiv 2023.07.17.549342 (2023) doi:10.1101/2023.07.17.549342.

40. Champer, J., Kim, I. K., Champer, S. E., Clark, A. G. & Messer, P. W. Suppression gene drive in continuous space can result in unstable persistence of both drive and wild-type alleles. Mol Ecol 30, 1086–1101 (2021).

41. Tolosana, I. et al. A Y chromosome-linked genome editor for efficient population suppression in the malaria vector *Anopheles gambiae*. bioRxiv 2024.05.14.594116 (2024) doi:10.1101/2024.05.14.594116.

